# A pan-genome approach to decipher variants in the highly complex tandem repeat of *LPA*

**DOI:** 10.1101/2022.06.08.495395

**Authors:** Chen-Shan Chin, Sairam Behera, Ginger A. Metcalf, Richard A. Gibbs, Eric Boerwinkle, Fritz J. Sedlazeck

## Abstract

Lp(a) is an important factor in Coronary Heart disease risk, and its expression is correlated with the length of the *LPA* gene. The KIV-2 region of *LPA* consists of variable tandem repeats whose number is highly variable across individuals. Little is known about the inner diversity of the KIV-2 repeats themselves. Here we utilize a pan-genome graph approach across 47 haplotype resolved assemblies to identify unexpected variation across KIV-2.

## Main

The Lp(a) protein and associated *LPA* gene have been extensively studied in the last 30 years. Interest stems from LP(a)’s demonstrated clinical utility as a predictor of vascular-related diseases^1^, its demonstrated heritability which has been estimated between 70-90%^2,3^, and the significant variability of plasma Lp(a) levels between race/ethnic groups^4–6^. Inter-individual differences in Lp(a) levels are extraordinary, spanning a range that is three-fold the mean ^7^. Likewise, individuals of African descent have 2 to 3 times higher Lp(a) plasma levels than European or Asian populations^8–11^. The preponderance of evidence suggests that these differences have genetic origins, but still no clear genomic explanation for these phenomena has been reported.

Lp(a) is encoded by the *LPA* gene (ENSG00000198670) on the reverse strand of chromosome 6 (6q27) of the human genome reference GRCh38. It consists of multiple kringle repeat units, which form a long gene body of 132,793 bp. *LPA* is hypothesized to have emerged from a recent gene duplication of *PLG* such that only humans, apes and old world monkeys carry the *LPA* gene^12,13,14^. Thus, *LPA* shares a high sequence similarity with *PLG* (78-100% identity)^7^. Expansion and differentiation in humans led to 10 different kringle domains that are all specific in their amino acid composition^7^. Depending on the species, certain kringle domains have been gained or lost. One kringle domain (kringle IV KIV) underwent a large expansion and is represented multiple times by KIV type 2 (KIV-2) whereas the other kringle IV domains are uniquely present (KIV-1, KIV3-10). The number of KIV-2 repeats have been observed to be inversely correlated with the expression of Lp(a), the levels of plasma Lp(a) levels and the risk of vascular diseases^10,15,16^ however the mechanism behind this correlation remains unknown. The KIV-2 repeat has been reported at approximately 5.5kbp long and encompasses two exons that are both expressed. KIV-2 is tandem repeated between 5 and 50+ times along the gene body itself^7^. KIV-2 includes two additional exons which, depending on the number of repeats, may make up 70% of the transcripted LPA mRNA length^17^.

Despite the importance, it remains difficult to determine the number of KIV-2 repeats directly from short or long read sequence data because of the general low mappability of sequencing reads across the *LPA* region on GRCh37 or GRCh38. Multiple studies have shown different SNV (e.g. rs3798220 and rs10455872) alleles that are within the *LPA* gene^3,18^, but outside of the KIV-2 repeat, to have an association with LP(a) levels and thus repeat number^7^. However, these SNVs only account for a minority of the variance in Lp(a) levels^19, 20^ and the results are race/ethnicity specific^21,22^. In addition, it has been suggested that the repeats (KIV-2) are under strong selection and therefore are nearly identical^7,13,38,39^.

Here we describe how long read data and a pan graph genome approach can be utilized to comprehensively identify and map the overall diversity of the *LPA* gene with a specific focus on the KIV-2 region. We show considerable diversity across its coding and noncoding region that was not observed previously, both within and among populations. We showcase the power of this approach by enabling alignment of short read sequence data back to the graph to characterize the haplotype frequency distribution across the region. These data not only represent important steps towards a deeper understanding of *LPA* diversity, but provide a low cost and scalable road map for characterizing other complex clinically relevant regions of the human genome^23–25^.

## Results

To first assess the *LPA* locus, we compared the newly released T2T assembly (CHM13)^26^ to the existing GRCh38^27^ reference. **Supplementary Figure 1A** shows an alignment of one of the assemblies (CHM13) to the GRCh38 reference genome showing a much higher number of KIV-2 repeats than represented in GRCh38 itself, which only represents six copies of KIV-2. The alignment underscores the previously reported high homology across KIV-2 repeat units, but also the ill representation of KIV-2 along GRCh38. Thus, we further compared the annotated KIV-2 sequence from Genebank (L14005.1) to the CHM13 reference sequence. We identified multiple subsegments that do not align well, indicating divergence from the Genbank sequence. Potentially more interesting is that the sequences (L14005.1 and CHM13) itself do not agree on the start (bottom of **Supplementary Figure 1B**) of the repeat regions. It actually only starts ∼3,800bp into the KIV-2 repeat. This is likely because Exon 1 was used as a target for the PCR primer for the annotation (L14005.1) of this region. Using the CHM13 as a reference point, we shifted the annotation of KIV-2 repeat such that it aligns with what we observe from the CHM13 assembly. We define the repeat array anchored repeat unit type I from chr6:161,982,999-161,988,542 and anchored repeat unit type II from chr6:161,911,055-161,916,599 in the CHM13 assembly. **Supplementary Figure 1C** shows the dot plot of the so modified start/stop of the KIV-2 repeat itself. This ∼3,800bp shift improves the overall alignment pattern as it shows that it fully contains the repeat itself (see methods). Noteworthy, is that this modification of start/stop of the KIV-2 repeat actually would shift the nomenclature of exon 1 and 2 since the original (Genebank L14005.1) started after exon 1.

To further explore the diversity of KIV-2 region, we obtained 47 Pacbio HiFi based long read assemblies that are phased and highly contiguous^28–30^. These 47 assemblies span a wide variety of samples from the United States and its territories (e.g. Puerto Rico) to Africa (e.g. Gambian Mandinka) to East Asia (e.g. China) and thus represent partially the diversity of *LPA* across multiple ethnicities. **Supplementary Table 1** gives details about the accessions and haplotypes that span *LPA* as five of these assemblies do not have both haplotypes represented across the *LPA* gene.This alignment step was done for all 47 assemblies (see **Supplementary material**). Given the evidence that these assemblies span the entire *LPA* locus, we next focused on the KIV-2 repeat motif.

**Figure 1A** shows a representation of the KIV-2 copy number repeats across individuals and haplotype. As previously reported, we observe highly different copy numbers of the KIV-2 repeats among individuals^20,31,32^. The y-axis of **Figure 1A** is sorted by ethnicity of the individuals. The colors in **Figure 1A** represent two diverse repeat KIV-2 units (yellow and magenta) that are 2.31% diverse (∼107 mismatches). Within the KIV-2 encoding repeat unit, the majority of the difference between the two types of repeats are concentrated in the short intron between the exons; a large portion of the mismatches are in the 1,549 bases between the exons. The blue segment represents a much further evolved unit (KIV-3) and marks the end of the KIV-2 repeats. We choose to include and label it as it shows that we reach the far end of the KIV-2 repeat units. Most of the time variable number of yellow repeats are followed by variable number of magenta repeats. Only in a few haplotypes of individuals (18.7%) the repeats are interspersed. There seem to be no clear association between the presence and number of magenta repeats and race/ethnicity.

**Figure 1:**
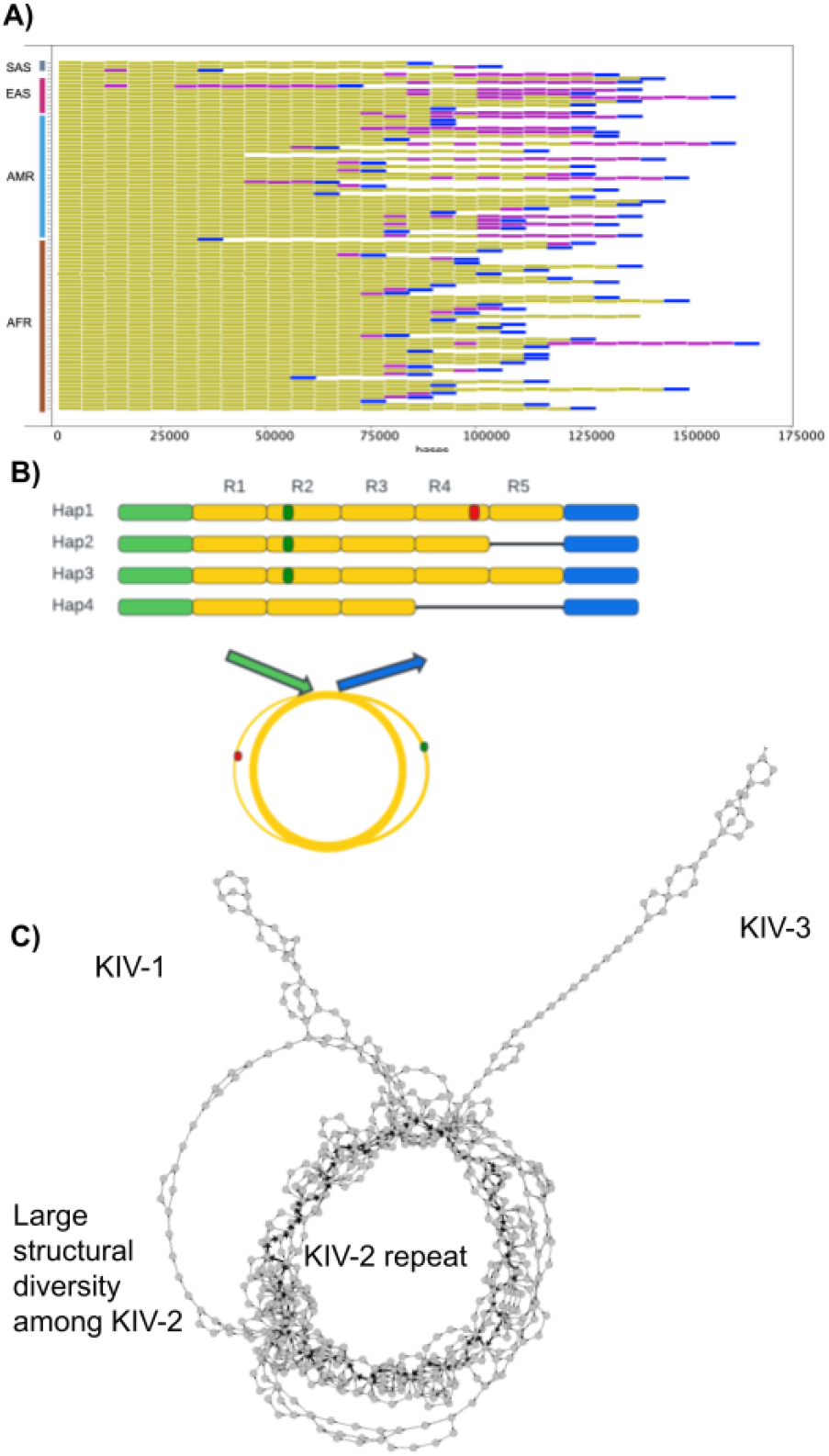
Insight into KIV-2 repeats. **A)** KIV-2 repeat differences across 47 assemblies (91 haplotypes) across multiple ethnicities. Repeats are labeled based on their sequence identity in yellow or magenta. Blue represents the KIV-3 unit and thus the end of the KIV-2 repeat cluster. (yellow: repeat with intron type 1, magenta repeat with intro type 2, blue, extend copy with part of KIV-3) **B)** Cartoon representation of the collapse of the repeats (R1-R6) across 4 haplotypes indicating SNV (common : green and rare : red). The thickness of the paths indicates the frequency of the alleles across multiple repeats. **C)** Graph genome representation of KIV-2 repeat region across the 47 haplotypes. The tandem duplication of the 5.5kbp unit is represented as a circularization where the sequence wraps around itself equal to the number of repeats per haplotype. We can observe remarkable diversity among the repeats indicated by the bubbles/loops that are branching out of the circle itself.

To further investigate the diversity across and within the repeats we utilized a graph/pan-genome approach. Graph Genome is a general data structure to encode multiple genomic sequences together as “graphs” to facilitate pan-genomics analyses^33–36^. This allows for a more wholesome representation across multiple individuals. Furthermore, advanced graph analysis techniques can help to analyze repeats and complex structures along the genome. To achieve this, we are using a minimizer approach to represent sequences in our graph as nodes (i.e., basic blocks). For each genomic sequence of interest, we first compute the minimizers specified by three parameters (*w*: window size, *k*: *k*-mer size and *r*: reduction factor)^37^ across the sequence. Thus, each genomic sequence (here from the assemblies) connects these minimizers together to form a path representing the individual region of here *LPA* KIV-2.

The advantage of this approach is that the parameters can be chosen to best represent certain variants. For example, for a large copy number of SV we would choose a large window and *k*-mer size. In contrast, if *w* and *r* parameters are chosen to be 1 the approach is equivalent to the de-Bruijn graph with a given *k*-mer size (*k*), and it will allow us to catch all variations. For the LPA repeat region, we use *w*=32, *k*=48, *r*=3. As we choose a small window and *k*-mer size to create dense minimizers across each DNA sequence, we can still find the correspondences from the short read sequencing data. This allows us to map the short read on to our graph by checking if a minimizer from the reads appears in the nodes. Furthermore, each of the edges is corresponding to a set of sequences between the two nodes, we can use those to relate a read into a set of nodes and edges in the graph for inferring the haplotypes. Thus, enabling a unique mapping and variant identification using short reads even in this repeat (KIV-2).

Applying this approach to encode the 47 long read assemblies (here 91 haplotypes) produces a graph representation of the 5.5 kbp KIV-2 repeat including its flanking region. **Figure 1B** shows a cartoon of this in a simplified manner. Here the individual repeats are collapsed in a ring structure with diversity indicated by deviations from the most supported (i.e., thickest) ring. **Figure 1C** shows the resulting graph structure across the 47 long read assemblies. The number of repeats is indicated by the number of times the circle is visited. The incoming and outgoing appendage in the graph on the top represents the start and the end of the KIV-2 repeats (i.e., KIV 1&3). **Figure 1C** shows that there are multiple alternative paths even with a stringent graph representation, indicating that the KIV-2 repeats are not as identical as previously postulated^31,32^. This also includes large alterations forming larger loops (e.g. left side). To further investigate these differences, we linearized this graph representation such that it only represents the 5.5kbp KIV-2 repeat unit which each individual passes multiple times.

**Figure 2A** shows a linear graph representation of the KIV-2 repeats along the 91 long read haplotypes. The width of the lines is corresponding to the number of times a path is supported by a single haplotype from the assembly. For **Figure 2A**, we use two red boxes to highlight the two exons coding regions along the graph (NM_005577.4). Overall **Figure 2A** allows us to clearly see the overall diversity of the intron between the KIV-2 two exons (highlighted in yellow). Other variants, such as the tandem repeat variation (highlighted in yellow) and other smaller deviations, are also evident.

**Figure 2:**
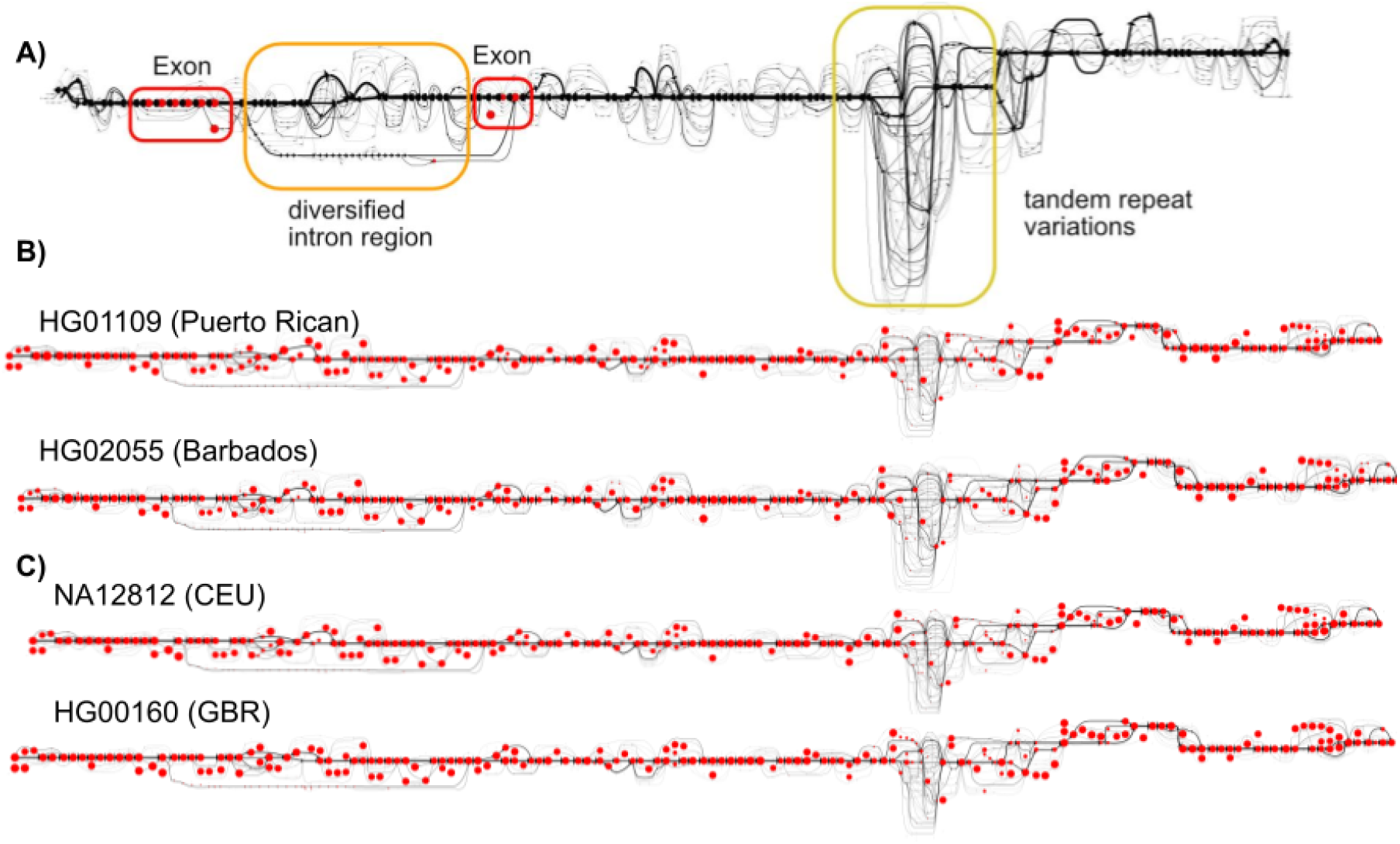
Linear graph genome representation of the 5.5kbp KIV-2 repeat unit. **A)** We identified a remarkable diversity across the KIV2 repeats within and among the 47 assemblies not only in the number of repeats but clearly indicating large gains and losses within the 5.5kbp repeats. This is shown by the different paths along the graph genome itself. **B)** This graph can be leveraged to map back short read data to KIV-2 directly. We show two samples (HG01109 and HG02055) whose data were included in the graph genome itself. Red circles indicate the path in the graph that was taken together with the width of the circle indicating the frequency of the reads. **C)** This graph representation also allows us to analyze other Illumina data across KIV-2 repeat. We show the mapping of two Illumina sequenced samples (NA12812 and HG00160) from European ethnicity. As before the red circles indicate the path in the graph that was taken together with the width of the circle indicating the frequency of the reads.

Next, we utilized this graph genome structure to not only map out the diversity but further enable the alignment of short reads. This would enable the assessment of *LPA* KIV-2 diversity at a much larger scale. **Figure 2B** shows the results of remapping two samples that are part of the initial set of assemblies. Here the black lines represent the individually observed haplotypes across one KIV-2 repeat. For each sample, we labeled each haplotype/path in the graph genome by its support. This is indicated by the red dots across each black line. The size of the dot reflects the number of Illumina reads that could be mapped to each path and does indicate the diversity that each sample includes. For example, HG01109 from Puerto Rican descent shows multiple paths throughout the graph itself. This is highlighted especially by multiple red dots indicating alternative paths over the KIV-2 repeat unit. Similarly, HG02055 from Barbados shows again a high diversity across the KIV-2 repeat unit. The samples also show diversity (i.e., alternative paths) across the two exon regions of the KIV-2 repeat unit. This clearly highlights not only the diversity of KIV-2 but also the utility of the graph genomes to improve the representation of variations across such a complex region itself. Maybe even more importantly it demonstrates that the graph can indeed be utilized to align short read data directly into the KIV-2 region to identify variation.

Lastly, we investigated if our graph genome approach further enables the analysis of other data sets to estimate e.g. allelic frequencies in the population. For this we aligned four European Illumina genome data (**Supplementary Figure 2**). **Figure 2C** shows two European samples along the same graph genome, which highlights the ability of our approach to further analyze samples that are initially not part of the graph genome itself. Again we found that even in these European samples multiple variants across KIV-2 region are present and that this region is far from homogenous.

## Conclusions

In this study we applied the concept of graph/pan-genomes to obtain insights into a single gene (*LPA*) that for decades buzzle the scientific community. Using a graph genome approach here not only shows to enable the mapping of short reads, but further allows us to identify yet undiscovered diversity among KIV-2 repeats within and among individuals. Thus, the suggestion that this locus is under strong selection and therefore the KIV-2 repeats are nearly identical^7,13,38,39^ seems to be invalid (**Figure 1C+2A-C**). The role of these mutations, however, remain for now unknown and might as well result in “null alleles” for Lp(a). What is needed is a more comprehensive approach utilizing long reads and RNA sequencing to further discover insights into the diversity of this important CHD risk gene. Nevertheless, what this study clearly shows is the importance and future role of graph based approaches not only to represent diversity but also to enable the utilization of existing or future short read data to study *LPA* and other medically important but challenging genes^23^.

## Methods

### Assembly comparison

**Supplementary Table 1** shows the source of the individual assemblies.

#### Identification of the repeat unit

We utilize the most recently assembled CHM13 T2T genome (v1.1 draft). We perform a self-self alignment of the sequence chm13_chr6:161762988-162032102 from CHM13 and identify the start point of the KIV-2 repeat array. We fetch the sequence from the starting point and identify the end of the repeat by identifying the same sequences from the starting point. The type I and type II template sequences we use for this study are extracted from the reverse complement of CHM13, chr6:161982999-161988542, and chr6:161911055-161916599, respectively.

#### Building of the graph

For **Figure 1**, a set of sparse minimizers^37,40^ of *k*=96, *w*=12, and *r*=2 are extracted and the graph is constructed as the genomic graph of the sequence of the minimizers across each contig. Each node in the mimizer-derived genomic graph is one of the mimizers and if a genomic contig has two consecutive mimizers in the sequence then an edge is added into the graph. Such mimizer-drived graphic graph allows us to capture the pangenomic structure with the level details showing how the KIV-2 repeat array relates to other parts of the gene. The repetitive sequences (hence the repetitive minimizer sequences) induced the circles shown in **Figure 1**.

For **Figure 2**, we are interested in the pangenomic structure that is more fine detail than those in **Figure 1** for a smaller region. We use *k*=48, *w*=32 and *r*=3 for the minimizer-derived genome graphs. As we use larger *k*-mer and small *r*, it allows us to capture more details of the repeat region even with a slightly larger *k*. It may be desirable to further reduce w and r to capture fine details for the difference. However, it may create graphs that may be hard to be comprehended by human eye.

#### Exon identification

We identify the exon sequences in KIV-2 with NM_005577.4 from GenBank (https://www.ncbi.nlm.nih.gov/nuccore/NM_005577.4). All 48 mers are extracted from the NM_005577.4 and compared to the kmer in the graph. The exact *k*-mer matches are used to identify the nodes in the graph that are corresponding the exon regions.

#### Remapping of short reads to the graph

Each of the 48-mer (unique within the repeat unit as we do have a loop in the graph) in the reads that are extracted As the graph node are unique in the graph. Such k-mers and the corresponded path derived from the reads can be easily identified as the nodes and edges in the graph and highlighted as the red nodes in **Figure 2**. We also count how many reads in each sample have the corresponded minimizers of the nodes. The sizes of the nodes in the graph is roughly proportional to the number of counts. The number of counts are determined by the copy number of the KIV-2 repeat and potential sequence variants.

## Data availability

**Supplementary Table 1** shows the source of the individual assemblies.

## Code availability

The library, data and the Jupyter Notebooks used to generate the plots are released as a Docker image that can be downloaded at https://hub.docker.com/r/cschin/pgr-lpa

## Acknowledgements

This study was partially supported by NIH grants (1U01HG011758-01).

