## Supplementary material for "A pan-genome approach to decipher variants in the highly complex tandem repeat of *LPA*"

1: Sema4, Stamford, CT 06902, USA

2: Human Genome Sequencing Center, Baylor College of Medicine, Houston, TX 77030, USA

3: Human Genetics Center, School of Public Health, University of Texas Health Science Center at Houston, TX 77030, USA

4: Department of Computer Science, Rice University, 6100 Main Street, Houston, TX 77005, USA

### Supplement

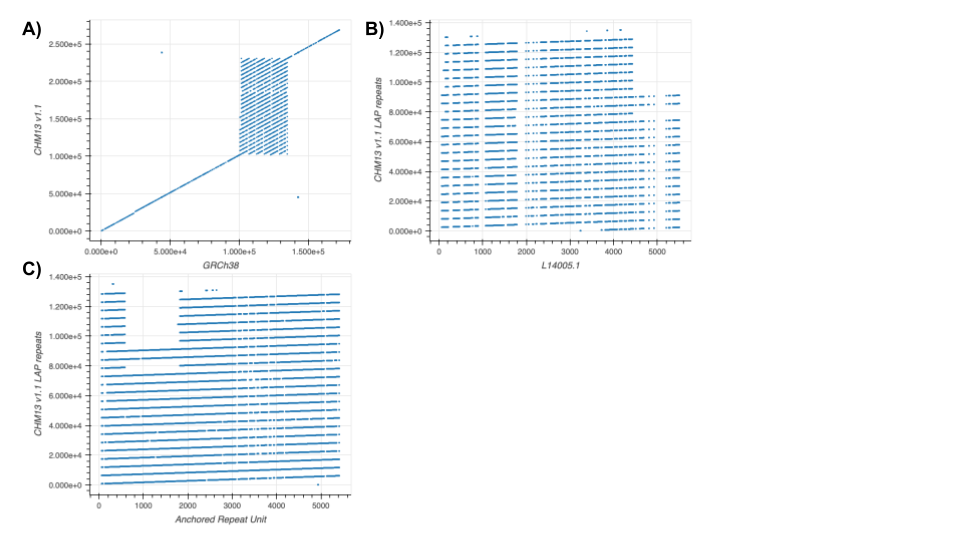

***Supplementary Figure 1:*** *Genome alignment representation of LPA.* ***A)*** *Visualization of the alignment of the CHM13 vs. GRCh38 assemblies of the LPA. Due to the repeat number difference, the typical alignment algorithm will break as there is no single one-to-one alignment across the repeat region. It may limit the variant-based analysis due to read mapping issues of the highly repetitive nature.* ***B)*** *Alignment comparison between reference genome CHM13 and the original KIV-2 repeat sequence (L14005.1). Of note is the lack of matching segments at the beginning of the regions in CHM13 (lower left of the alignment).* ***C)*** *Alignment comparison of a shifted CHM13 KIV-2 repeat sequence proposed here to better align with the reference CHM13 genome. This shifted start of KIV-2 repeat causes the sequence to start with Exon 2 from L14005.1*

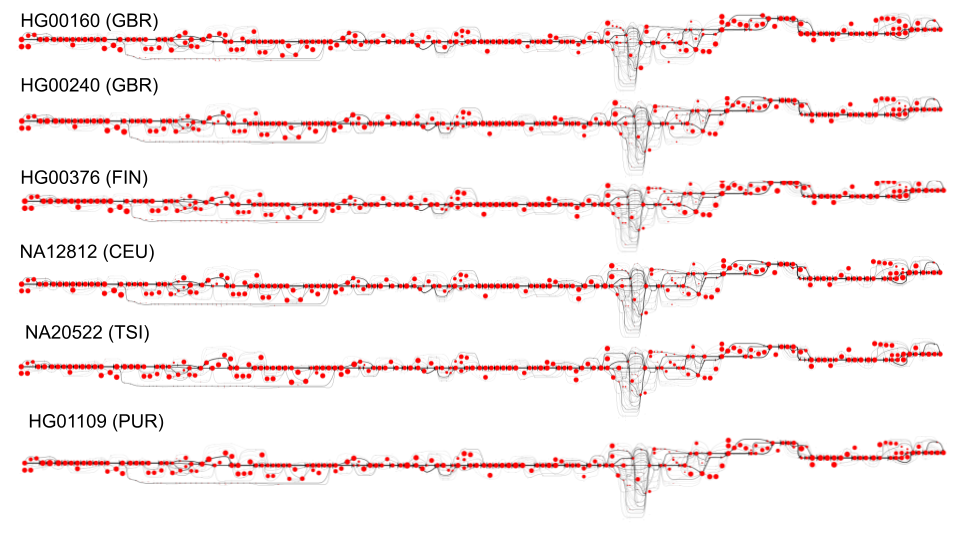

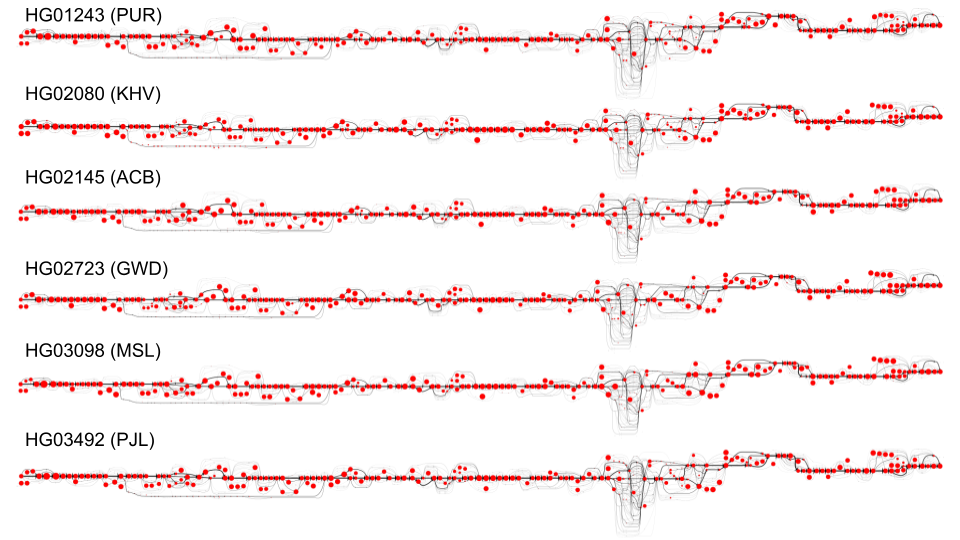

***Supplementary Figure 2:*** *Mapping of illumina short read data from various ethnicities to KIV-2 region of graph genome.*
